# Comparative evaluation of the genomes of common bacterial members of the *Drosophila* intestinal community

**DOI:** 10.1101/036020

**Authors:** Kristina Petkau, David Fast, Aashna Duggal, Edan Foley

## Abstract

*Drosophila melanogaster* is an excellent model to explore the molecular exchanges that occur between an animal intestine and their microbial passengers. For example, groundbreaking studies in flies uncovered a sophisticated web of host responses to intestinal bacteria. The outcomes of these responses define critical events in the host, such as the establishment of immune responses, access to nutrients, and the rate of larval development. Despite our steady march towards illuminating the host machinery that responds to bacterial presence in the gut, we know remarkably little about the microbial products that influence bacterial association with a fly host. To address this deficiency, we sequenced and characterized the genomes of three common *Drosophila*-associated microbes: *Lactobacillus plantarum, Lactobacillus brevis* and *Acetobacter pasteurianus*. In each case, we compared the genomes of *Drosophila*-associated strains to the genomes of strains isolated from alternative sources. This approach allowed us to identify molecular functions common to *Drosophila*-associated microbes, and, in the case of *A. pasteurianus*, to identify genes that are essential for association with the host. Of note, many of the gene products unique to fly-associated strains have established roles in the stabilization of host-microbe interactions. We believe that these data provide a valuable starting point for a more thorough examination of the microbial perspective on host-microbe relationships.

## INTRODUCTION

Environmental, microbial, and host factors act at mucosal barriers to establish a unique microclimate that shapes the lives of all participant species (1). For example, expression of a host genotype in gastrointestinal tissues works in concert with extrinsic factors to determine microbial associations (2). For their part, the metabolic outputs of the gastrointestinal microflora influence critical events in the host such as education of immune phenotypes (3, 4), development of intestinal structures (5), and access to essential micronutrients (6). Given the intertwined relationship between host phenotype and microbial genotype, it is of some surprise that hosts often tolerate extensive alterations to their microflora in response to environmental shifts such as changes in diet (7). However, alterations to the gastrointestinal microflora are not invariably without consequence, and intestinal dysbiosis may lead to chronic, debilitating, and occasionally deadly diseases within the host (8–12). Our appreciation of the holobiont as an intricate network of biochemical and genetic transactions between multiple participants mandates a thorough evaluation of the microbial genomes that shape host physiology. Unfortunately, such studies are tremendously complex in conventional mammalian models due to the size of the microbiome, and the lack of laboratory techniques for the isolation and manipulation of most intestinal microbes.

The simple invertebrate *Drosophila melanogaster* is an excellent model holobiont (13, 14). From a developmental perspective, the *Drosophila* posterior midgut shares a number of important similarities with the small intestine of more complex mammalian counterparts (15). Both organs are endodermal in origin, and are surrounded by a sheath of mesodermal visceral muscle (16, 17). The mammalian small intestine and *Drosophila* posterior midgut are maintained by regularly spaced, basal intestinal stem cells that generate transitory progenitor cells (18–20); the non-proliferative enteroblasts of *Drosophila*, and the transient-amplifying cells of mammals. In both systems, signals along the Notch-Delta axis promote differentiation of transitory progenitors into secretory enteroendocrine cells or absorptive enterocytes (15, 21). In contrast to mammals, *Drosophila* lacks specialized basal paneth cells for the release of antimicrobial peptides. Nonetheless, the fly genome encodes antimicrobial peptides that actively contribute to the control of intestinal symbionts and pathogens (22), suggesting release of such factors into the *Drosophila* intestinal lumen. In both the mammalian small intestine and *Drosophila* midgut, host factors and biogeography favor association with members of the *Lactobacillaceae* family (2, 23). In return, metabolites from *Lactobacilli* activate host response pathways that promote intestinal stem cell proliferation (24). Combined with the genetic accessibility of flies and their suitability for longitudinal studies of large populations in carefully defined environments, *Drosophila* is an excellent system to decipher the forces that determine genetic interactions within a holobiont (13, 25, 26).

In contrast to conventional vertebrate models, the *Drosophila* microbiome consists of a small number of bacterial species that are easily isolated and cultured (27). The adult *Drosophila* intestine hosts little to no bacteria immediately after emergence from the pupal case, and the microfloral population grows in number over time (28). Several studies established that environmental factors and host genotype influence the diversity of the microflora (22, 29, 30). It is unclear if bacteria establish stable associations with the host gut, or if they cycle from the intestine to the environment and back (31). Nonetheless, lab-raised or wild *Drosophila* frequently associates with representatives of the genii *Lactobacillus* and *Acetobacter*. These data suggest that the intestinal lumen of an adult fly favors the survival of specific bacteria, and that such bacteria encode the necessary factors to survive or proliferate within a *Drosophila* intestine.

Consistent with a long-term association between the fly intestine and specific microbes, many *Drosophila* phenotypes are influenced by individual *Lactobacillus* or *Acetobacter* species. For example, *Lactobacillus plantarum*, a common *Drosophila*-associated microbe, promotes larval development via regulation of the TOR signal transduction pathway (32), while *Acetobacter pomorum* regulates host Insulin Growth Factor signals to promote development and metabolic homeostasis (33). In addition, members of the *Acetobacter* and *Lactobacillus* populations regulate levels of essential nutrients in the host (34–36). Combined, these data present a compelling argument that *Lactobacilli* and *Acetobacter* are important members of the *Drosophila*-microbe holobiont.

Despite our advances in the elucidation of *Lactobacillus* and *Acetobacter* influences on their *Drosophila* host, we know remarkably little about the bacterial genomic elements that determine microbial responses to, and survival within, the *Drosophila* adult intestine. To address this issue, we prepared whole genome sequences of three bacterial species that regularly associate with *Drosophila* – *Lactobacillus brevis, Lactobacillus plantarum*, and *Acetobacter pasteurianus*. These sequences included those for a *Lactobacillus plantarum* strain isolated from our lab-raised flies, and a separate strain isolated from a wild *Drosophila*. For each species, we compared *Drosophila*-associated bacterial genomes, including ones reported previously, to the genomes of reference strains isolated from non-*Drosophila* sources. We focused on biochemical and signal transduction pathways that respond to environmental factors, as we reasoned that *Drosophila*-associated genomes will display signatures of adaptation to association with their fly host. Our studies identify several biochemical pathways that are enriched in the genomes of *Drosophila*-associated *Lactobacilli*, and uncover *Acetobacter* genomic elements that regulate viability within the host.

## RESULTS

Intestinal bacteria face structural and chemical barriers that typically deter association with an animal intestine. In response, microbial genomes encode factors that overcome host defenses to permit bacterial growth or survival. Here, we examined the genomes of *L. brevis, L. plantarum* and *A. pasteurianus*, common members of the *Drosophila* intestinal community. For each species, we studied whole-genome sequences of bacterial strains isolated from *Drosophila* intestines, and compared them to related strains isolated from the environment. As it is unclear if *Lactobacilli* or *Acetobacter* establish autochtonous or allochtonous relationships with the adult intestine we will refrain from use of the term commensal throughout this study. Instead, we will identify bacterial strains isolated from flies as associated with *Drosophila*, and we will identify all other strains as environmental isolates. Details on the respective genomes characterized in this study are presented in table 1.

**Table 1:** Details on the eleven bacterial strains used in this study.

| Bacteria | Strain | Isolated from<br>*Drosophila* | Reference |
| --- | --- | --- | --- |
| *Lactobacillus brevis* | ATCC 367 | No | (72) |
| *Lactobacillus brevis* | EW | Yes | (37) |
| *Lactobacillus brevis* | EF | Yes | This study |
| *Acetobacter pasteurianus* | NBRC 101655 | No | (73) |
| *Acetobacter pasteurianus* | ATCC 33445 | No | (73) |
| *Acetobacter pasteurianus* | AD | Yes | This study |
| *Lactobacillus plantarum* | JOJTO1.1 | No | (49) |
| *Lactobacillus plantarum* | ATCC 14917 | No |  |
| *Lactobacillus plantarum* | WJL | Yes | (48) |
| *Lactobacillus plantarum* | KP | Yes | This study |
| *Lactobacillus plantarum* | DF | Yes | This study |

We processed each genome in a similar manner. Where necessary, we used genomic databases to identify the bacterial species of newly sequenced genomes. We then annotated each genome with RAST, used PHAST to scan each genome for intact prophages, and searched for possible CRISPR arrays in the respective genomes. We scrutinized the annotated genomes for functions that facilitate microbial survival within the intestinal lumen, with a focus on genes involved in signal transduction, transcriptional responses, orchestration of stress responses, or induction of virulence factors. Finally, we compared environmental and *Drosophila*-associated genomes for each species to identify bacterial factors that are unique to *Drosophila*-associated genomes. We present the results for each species below.

### Lactobacillus brevis

### General Genomic Features

*L. brevis* frequently associates with *Drosophila* intestines, and the whole genome sequence of a fly-associated strain, *L. brevis* EW is available (37). We prepared a whole-genome sequence of an additional *L. brevis* strain (*L. brevis* EF) that we isolated from the intestines of wild-type adult *Drosophila* in our lab. For comparative purposes, we extended our study to include the genome of the environmental ATCC 367 strain. We found that *L. brevis* ATCC 367 readily associated with wild-type adult *Drosophila*. Genome-to-genome distance calculations suggest that the *Drosophila*-associated EW and EF strains are more closely related to each other than to the ATCC 367 strain (Table 2). The genomes of *Drosophila*-associated strains are also larger than the environmental strain, with approximately 500,000 nucleotides more, and an additional 500 coding sequences (Table 2, and Figure 1A).

**Figure 1:**
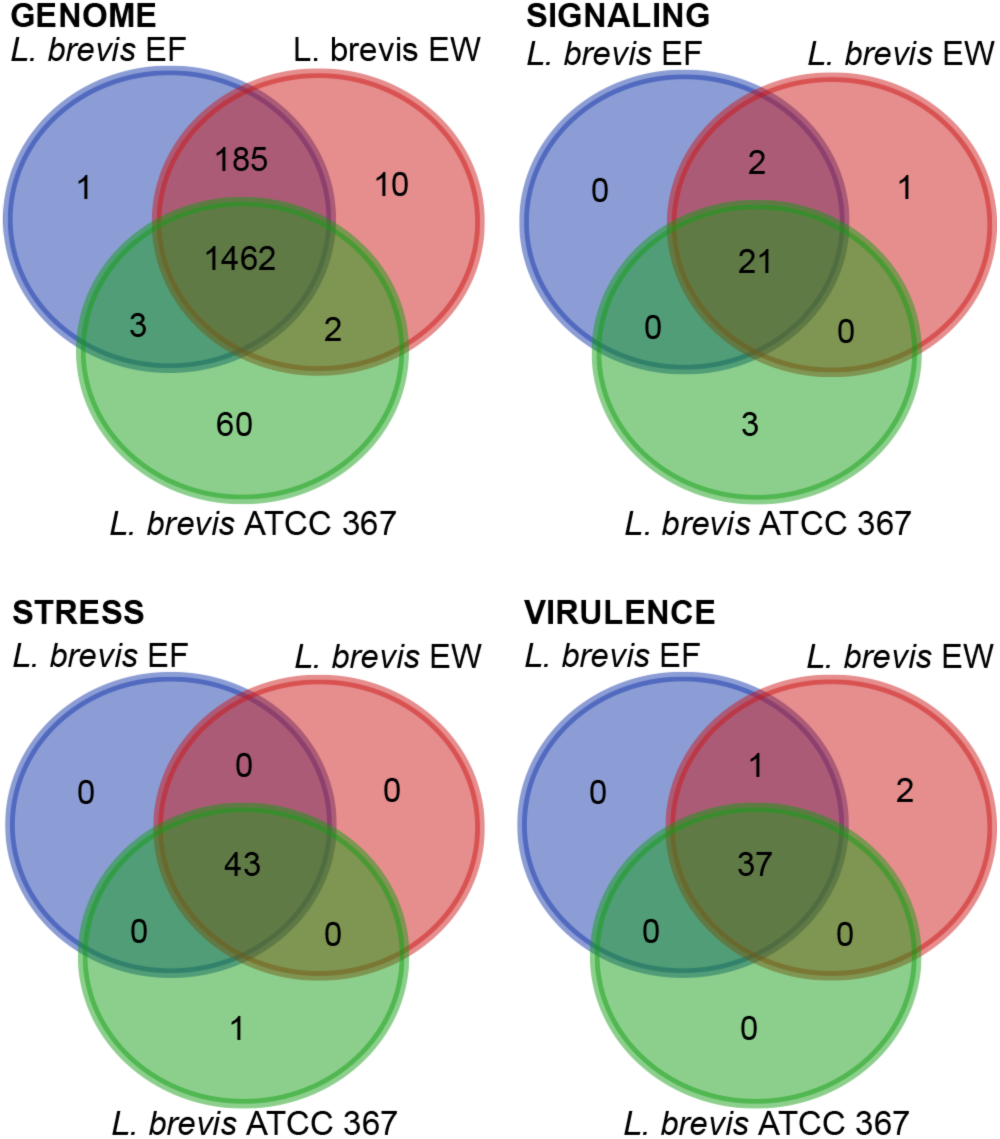
Distribution of unique gene functions in the genomes of *L. brevis* strains EW, EF and ATCC 367. All data are based on gene function annotations within RAST and exclude gene products with unknown functions.

**Table 2:** Details on *Lactobacillus brevis* genomes described in this study

|  | *L. brevis* EF | *L. brevis* EW | *L. brevis* ATCC #367 |
| --- | --- | --- | --- |
| Number of contigs or scaffolds | 32 | 38 | 3 |
| Genome to genome distance (Prob. DDH >= 70%) | 100 | 98.3 | 89.92 |
| Genome size | 2,864,530 | 2,885,101 | 2,340,228 |
| CG content (%) | 45.3 | 45.3 | 46.1 |
| Number of RNAs | 80 | 82 | 79 |
| Predicted CDS | 2808 | 2830 | 2284 |
| Assigned function | 1961 | 1969 | 1786 |
| Uncharacterized | 13 | 13 | 8 |
| Conserved hypothetical | 11 | 9 | 5 |
| Unknown function | 31 | 31 | 32 |
| Hypothetical | 673 | 682 | 431 |
| Phage-associated proteins | 119 | 126 | 22 |

#### Environmental Response Factors

We then examined genetic regulatory networks within the individual *L. brevis* strains. We hypothesized that *Drosophila*-associated strains will encode distinct regulatory components adapted to the harsh environment of an adult intestine. For these studies, we paid particular attention to two-component systems, transcription factors and additional DNA-binding proteins within the respective genomes. To our surprise, we did not observe substantial differences between *Drosophila*-associated and environmental genomes (Table 3, and supplementary tables 1–3), with the exception of a greater number of transcription factors in the *Drosophila*-associated genomes. Likewise, we only observed slight differences between *Drosophila*-associated and environmental strains when we considered genes dedicated to signal transduction, stress responses, or virulence (Figure 1B-D and supplementary tables 4–6).

**Table 3:**
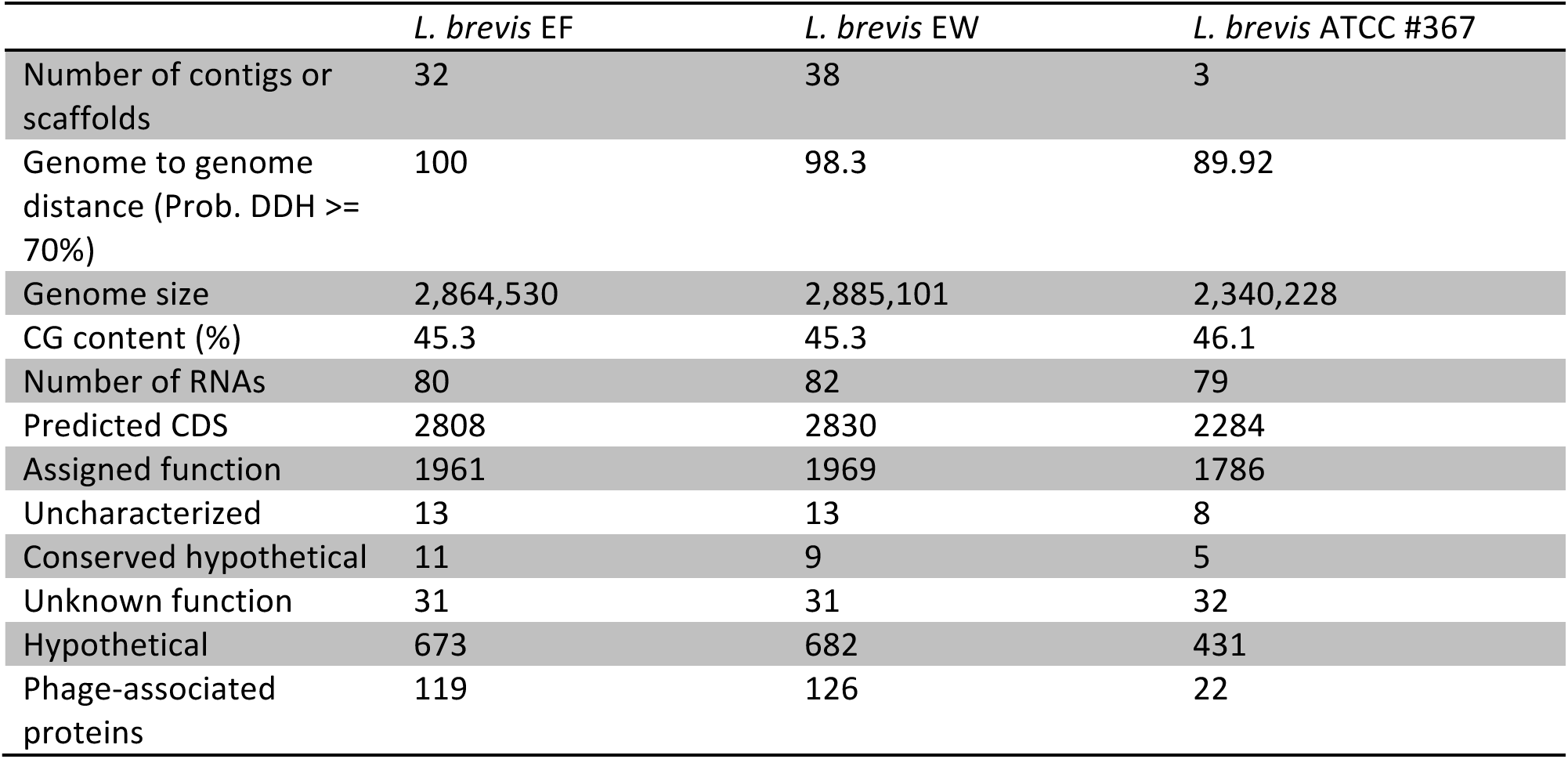
Organization of regulatory proteins within *Lactobacillus brevis* genomes described in this study

#### Prophages and CRISPR Responses

In contrast to the similarities between environmental and *Drosophila* associated strains noted above, we detected large differences when we searched for prophages within the respective genomes. The EW and EF genomes include four intact temperate prophages, compared to an absence of prophages from the environmental strain genome (Table 4). We found CRISPR sequences that target a common *Lactobacillus* phage within all three genomes, while the environmental strain encoded a separate CRISPR array that targets a *Lactobacillus* plasmid (Table 4). These results suggest an ongoing interaction between prophages and CRISPR defenses in the genomes of *Drosophila*-associated *L. brevis* strains.

**Table 4:** Prophage and CRISPR identification in *Lactobacillus brevis* genomes described in this study

|  | *L. brevis* EF | *L. brevis* EW | *L. brevis* ATCC #367 |
| --- | --- | --- | --- |
| Number of Prophages | 4 | 4 | 0 |
| Identity of Prophages | 3 X NC_019489<br>(*Lactobacillus* phage Sha1) | 3 X NC_019489<br>(*Lactobacillus* phage Sha1) |  |
|  | NC_027990<br>(*Lactobacillus* phage LBR48) | NC_009813 (*Listeria* phage B054) |  |
| Number of CRISPRs | 1 | 1 | 2 |
| CRISPR targets | *Lactobacillus* phage LBR48 | *Lactobacillus* phage LBR48 | *Lactobacillus* plasmid sequences |
|  |  |  | *Lactobacillus* phage LBR48 |

#### Function-Based Comparisons of Commensal and Reference Strains of *Lactobacillus brevis*

The data above point to subtle differences between the genomes of *Drosophila*-associated and environmental strains of *L. brevis*. To characterize these differences in greater detail, we performed a function-based comparison of the 185 genes that are common to *Drosophila*-associated genomes, but absent from the environmental strain. This set of 185 genes describes thirteen distinct functional categories, with forty-seven unique roles (Table 5). Perhaps unsurprisingly, phage and CRISPR-associated gene products account for two of those categories, and cover eleven of the forty-seven unique roles.

**Table 5:**
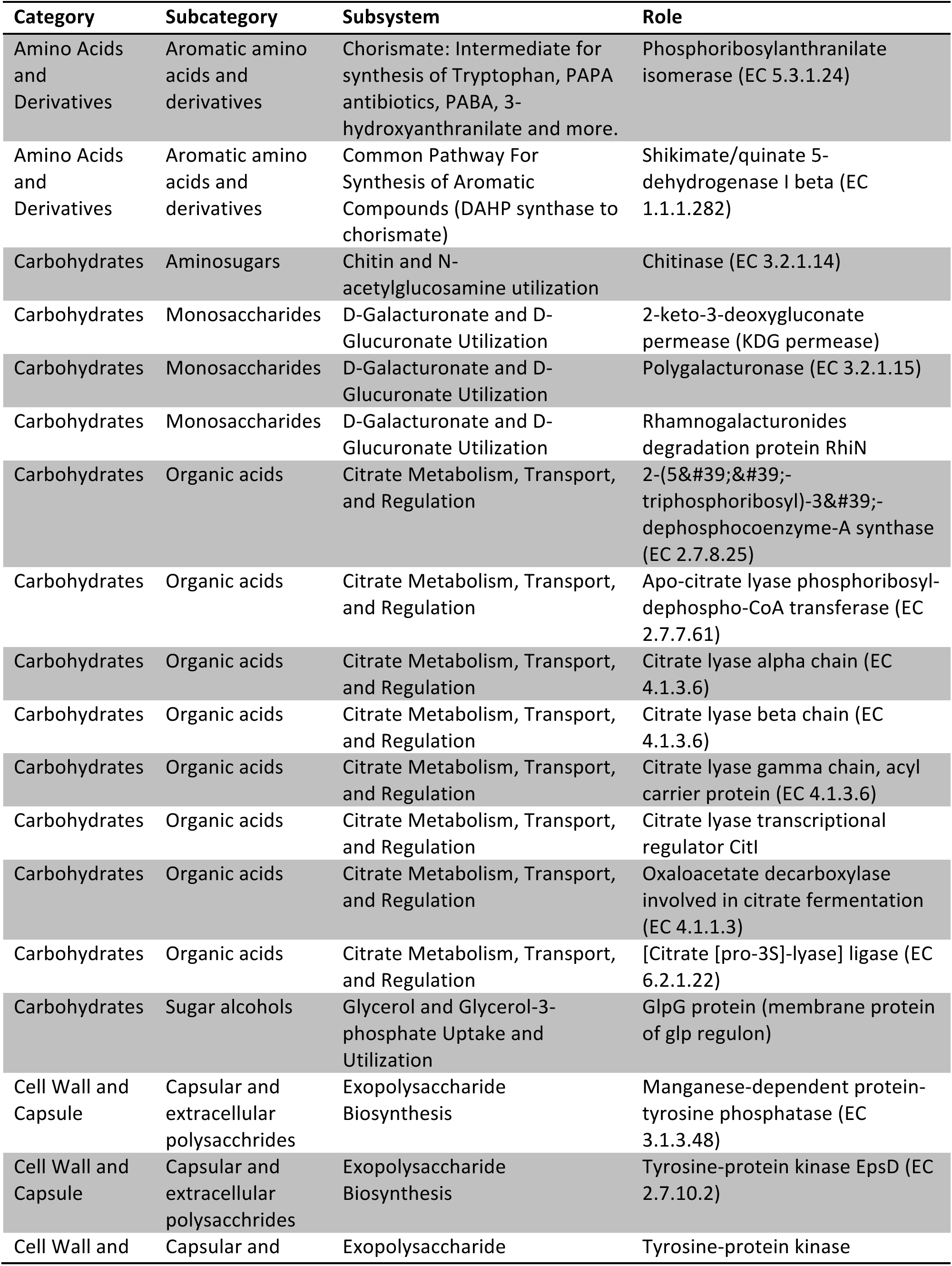
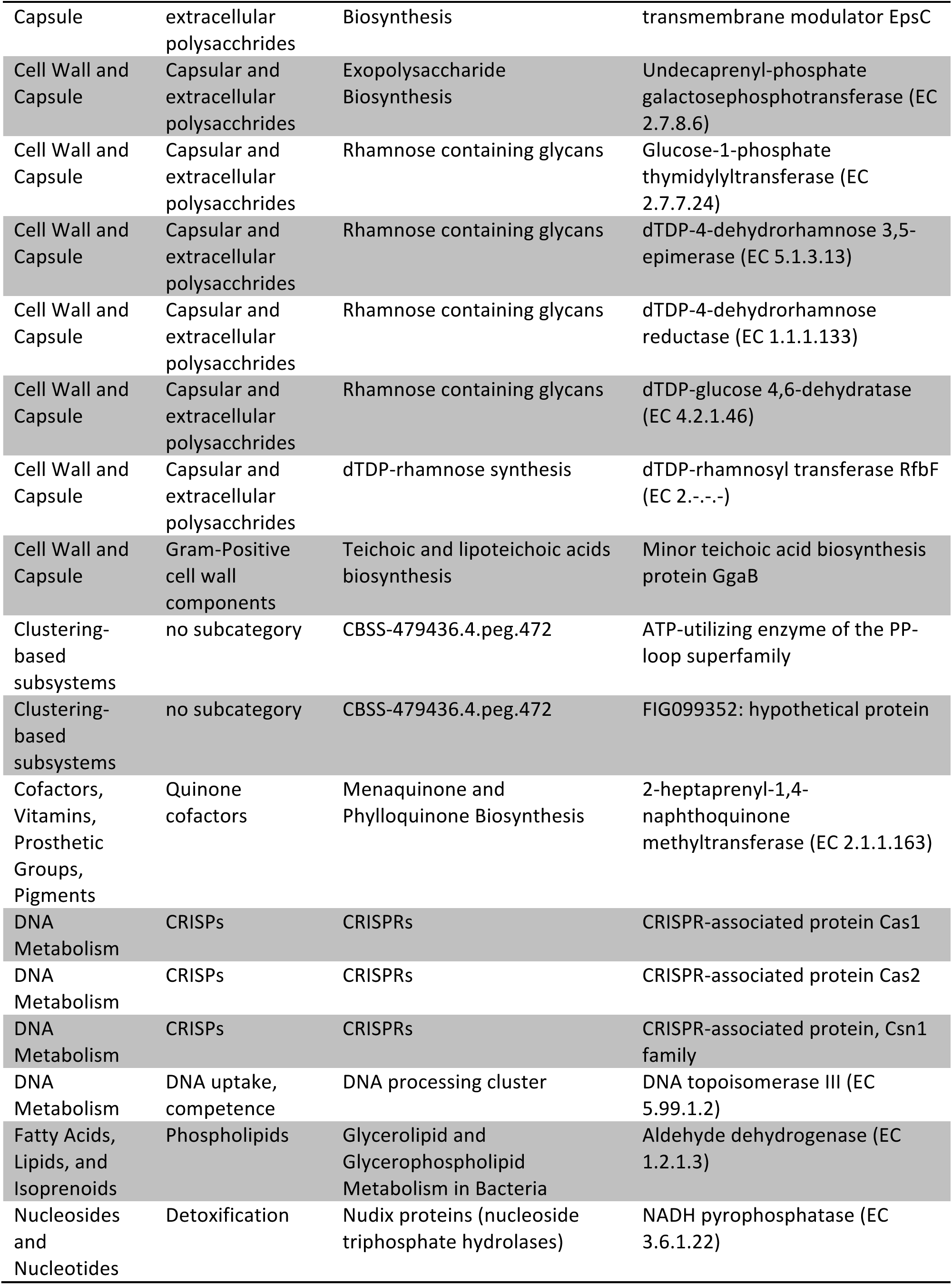
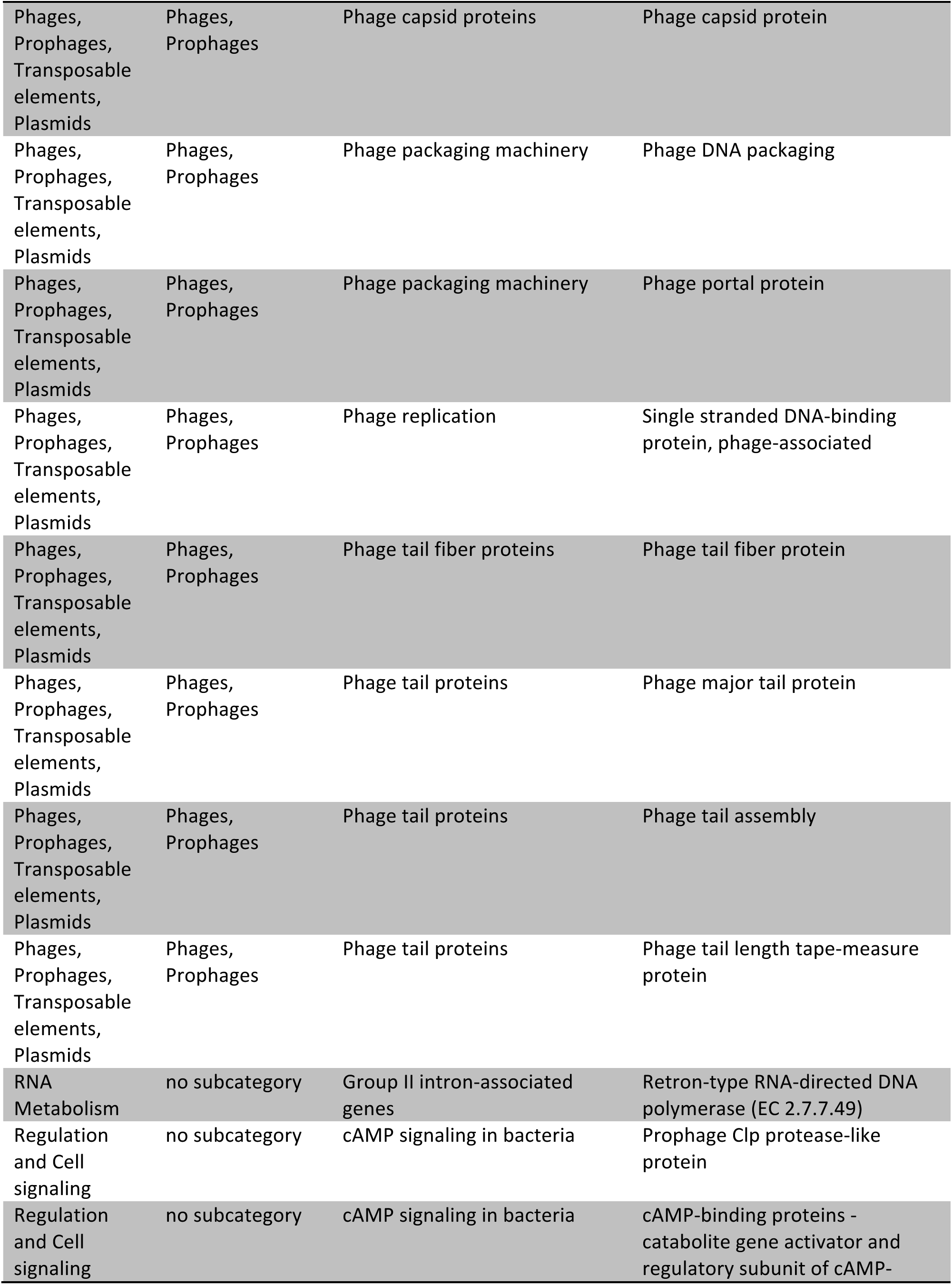
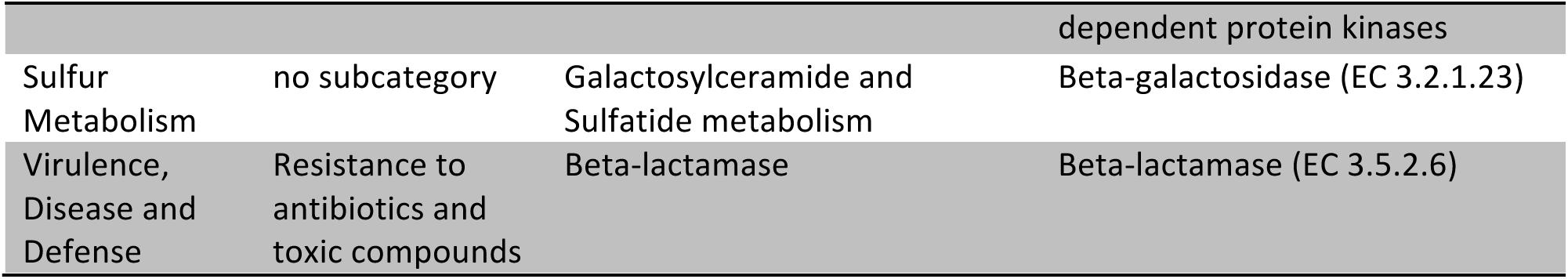
Function-based identification of systems unique to commensal strains of *Lactobacillus brevis*.

Of the remaining gene products, the dominant functional categories are dedicated to roles suited for survival within an intestine. For example, both *Drosophila*-associated genomes encode a biochemical cascade that converts α-D-glucose-1-phosphate to dTDP-4-dehydro-6-deoxy-L-mannose, an exopolysaccharide that contributes to biofilm formation, and prokaryotic survival within a host intestine (38, 39). *Drosophila*-associated strains of *L. brevis* also encode gene products that contribute to the formation of rhamnose-containing glycans, a cell membrane component of acid-fast bacteria that affects host-microbe interactions, such as adhesion, recognition, and biofilm formation (40).

We also identified gene products within *Drosophila*-associated genomes that facilitate nutrient acquisition from different sources (Table 5). Bacteria frequently respond to limitations in nutritional environments through activation of the cAMP Receptor Protein, a transcription factor that we did not identify in the environmental strain of *L. brevis*, but found in both *Drosophila-associated* strains. The cAMP Receptor Protein controls, among other things, the expression of gene products that coordinate metabolism of citrate (41), a function that is also enriched among associated *Drosophila*-associated *L. brevis* genomes. In lactic acid bacteria, citrate lyase is activated in acidic environments such as those found in the gut, and increases carbon utilization and energy generation by blocking the inhibitory effects of the *Lactobacillus* fermentation product lactate (42). Finally, we detected an enrichment of genes involved in the transport and degradation of pectin in *Drosophila*-associated *L. brevis* genomes. Pectin is an abundant source of energy and carbon for bacteria that grow on plant and vegetable surfaces, and microbial consumption of pectin accelerates the decay of organic matter. As *Drosophila* feed on bacteria and yeast that grow on decomposing fruit, we believe that pectin metabolism by the EW and EF strains of *L. brevis* increases the likelihood of their association with *Drosophila* in the wild.

### Acetobacter pasteurianus

### General Genomic Features

Although *Acetobacter* frequently associate with the intestines of *Drosophila* in the wild and in the lab, we are unaware of any whole-genome sequences of *A. pasteurianus* strains derived from the intestines of adult *Drosophila*. We consider this a significant deficit given the impacts of *Acetobacter* on *Drosophila* development and metabolism. To address this shortcoming, we completed a whole-genome sequence of an *A. pasteurianus* strain (A. *pasteurianus* AD) that we isolated from the intestines of wild-type *Drosophila*. For comparative purposes, we examined the available genomic sequences of the NBRC 101655 strain, and the ATCC 33445 strain. The NBRC 101655 strain was originally isolated from pineapple in Thailand and successfully associates with adult *Drosophila* (Figure 2). In contrast, we found that the environmental ATCC 33445 isolate fails to survive within the intestine of adult *Drosophila* (Figure 2).

**Figure 2:**
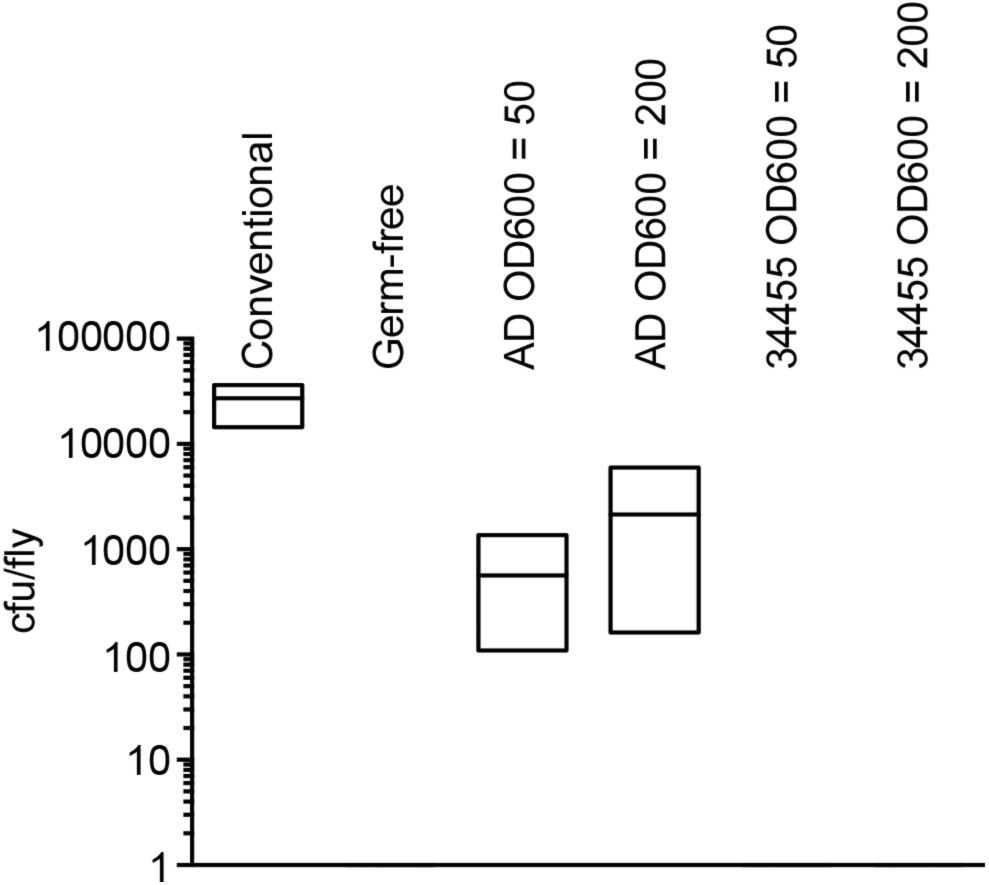
Quantification of *A. pasteurianus* association with conventionally reared (column 1) flies, germ-free (column 2) flies, gnotobiotic flies that were fed *A. pasteurianus* strain AD at OD600 of 50 and 200, respectively (columns 3 and 4), or gnotobiotic flies that were fed *A. pasteurianus* strain ATCC 33445 at OD600 of 50 and 200, respectively (columns 5 and 6). Each column shows the results of three separate measurements, and association was measure as bacterial colony-forming units per fly.

At first glance, we did not observe substantial differences between the respective genomes. Each genome is approximately 3 MB in length, with similar GC content, and similar numbers of RNA, and predicted coding sequences (Table 6). From an evolutionary perspective, *A. pasteurianus* AD appears more closely related to the NBRC 101655 strain than the ATCC 33445 strain (Table 6). Consistent with a greater evolutionary distance to the ATCC strain, we found that the ATCC 33445 genome encodes 112 unique proteins, while the NBRC 101655 and AD strains share 112 genes that are absent from the ATCC 33445 genome (Figure 3A).

**Table 6:** Details on *Acetobacter pasteurianus* genomes described in this study

|  | *A. pasteurianus* AD | *A. pasteurianus* ATCC 33445 | *A. pasteurianus* NBRC 101655 |
| --- | --- | --- | --- |
| Number of contigs or scaffolds | 161 | 306 | 294 |
| Genome to genome distance (Prob. DDH >= 70%) | 100 | 90.67 | 98.3 |
| Genome size | 2,830,055 | 2,888,200 | 3,018,312 |
| CG content (%) | 52.7 | 53.1 | 52.7 |
| Number of RNAs | 42 | 44 | 42 |
| Predicted CDS | 2673 | 2797 | 2834 |
| Assigned function | 1845 | 1866 | 1923 |
| Uncharacterized | 10 | 9 | 10 |
| Conserved hypothetical | 8 | 9 | 8 |
| Unknown function | 12 | 15 | 11 |
| Hypothetical | 770 | 854 | 852 |
| Phage-associated proteins | 28 | 44 | 30 |

**Figure 3:**
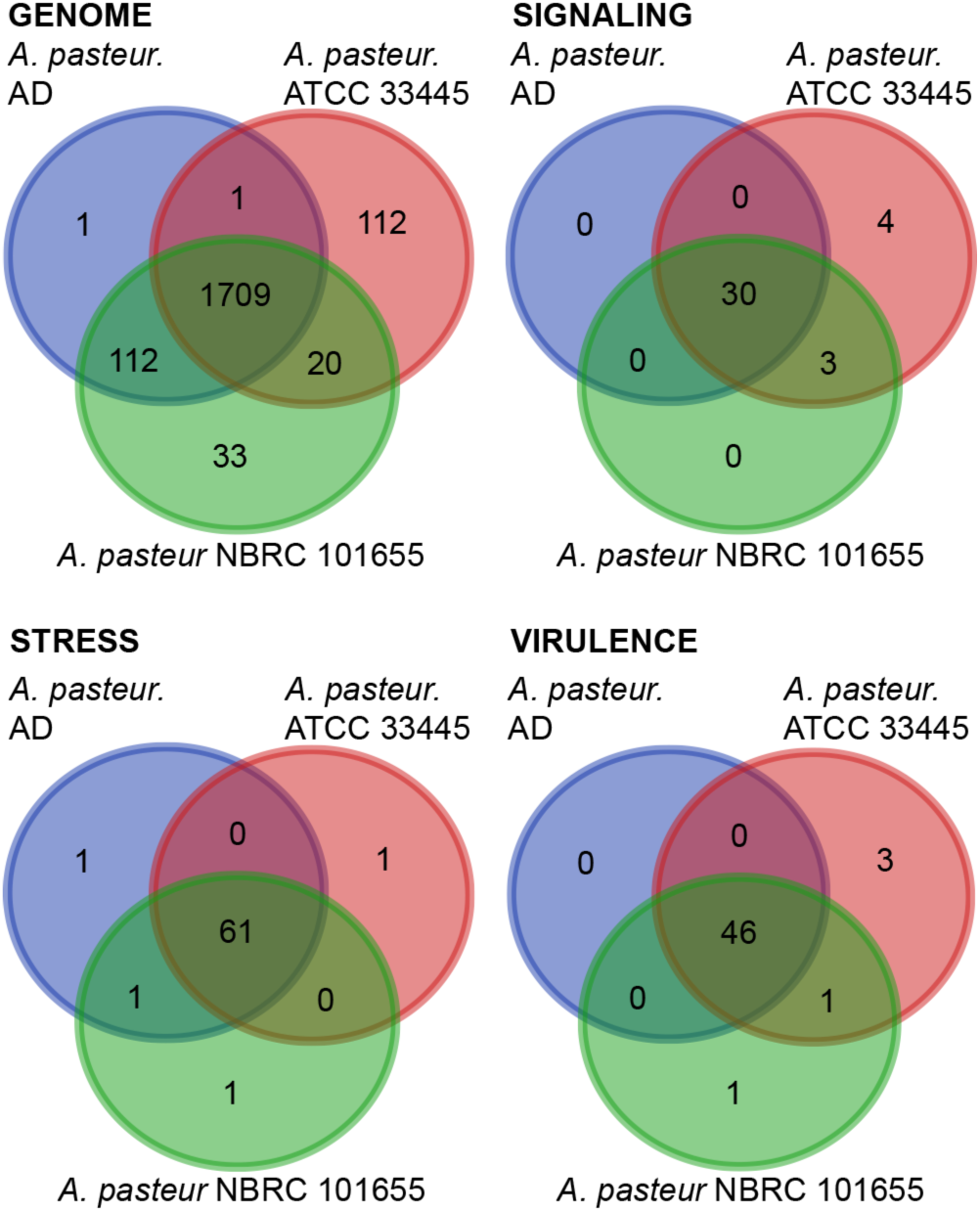
Distribution of unique gene functions in the genomes of *A. pasteurianus* strains AD, ATCC 33445 and NBRC 101655. All data are based on gene function annotations within RAST and exclude gene products with unknown functions.

#### Environmental Response Factors

As with *L. brevis*, we first compared the *Drosophila*-associated and environmental genomes for distinctions in gene products that control responses to environmental factors. Specifically, we looked at signaling factors (Figure 3B, supplementary table 7), stress response factors (Figure 3C, supplementary table 8), and virulence factors (Figure 3D, supplementary table 9). We also identified two-component systems, transcription factors, and other DNA binding factors in each genome (Table 7, and supplementary tables 10–12). Across this series of comparisons, the most pronounced differences were commensurate with a closer relationship of the AD strain to the NBRC 101655 strain than to the ATCC 33445 strain. Thus, this admittedly limited comparison does not appear to identify genomic components that readily distinguish *Drosophila*-associated *A. pasteurianus* genomes from environmental counterparts. Nonetheless, these functional characterizations uncover differences between the AD and NBRC 101655 *A. pasteurianus* genomes that associate with *Drosophila*, and the ATCC 33445 genome that fails to do so.

**Table 7:** Organization of regulatory proteins within *Acetobacter pasteurianus* genomes described in this study

|  | *A. pasteurianus* AD | *A. pasteurianus* ATCC 33445 | *A. pasteurianus* NBRC 101655 |
| --- | --- | --- | --- |
| Two-component systems | 28 | 26 | 28 |
| Transcription Factors | 105 | 100 | 127 |
| Other DNA binding proteins | 19 | 15 | 20 |

#### Function-Based Comparisons of Commensal and Reference Strains of *Acetobacter pasteurianus*

Our fortuitous identification of an environmental *A. pasteurianus* strain that fails to survive passage through the fly gut allowed us to explore *A. pasteurianus* genomes for factors that permit association with *Drosophila*. As the AD and NBRC 101655 strains successfully pass through the adult intestine, we reasoned that the AD and NBRC 101655 genomes encode biochemical functions absent from ATCC 33445 that permit association with *Drosophila*, or that the ATCC 33445 genome encodes biochemical functions absent from the other strains that prevent association with *Drosophila*. This prompted us to identify biological subsystems shared exclusively by the AD and NBRC 101655 (Table 9), or unique to the ATCC 33445 genome (Table 10).

**Table 8:** Prophage and CRISPR identification in *Acetobacter pasteurianus* genomes described in this study

|  | *A. pasteurianus* AD | *A. pasteurianus* ATCC 33445 | *A. pasteurianus* NBRC 101655 |
| --- | --- | --- | --- |
| Number of Prophages | 0 | 0 | 0 |
| Number of CRISPRs | 0 | 0 | 0 |

**Table 9:** Identification of RAST Subsystems absent from the ATCC 33445 strain of *Acetobacter pasteurianus*.

| Category | Subcategory | Subsystem | Role |
| --- | --- | --- | --- |
| Amino Acids and Derivatives | Alanine, serine, and glycine | Glycine Biosynthesis | Low-specificity L-threonine aldolase (EC 4.1.2.5) |
| Amino Acids and Derivatives | Arginine; urea cycle, polyamines | Polyamine Metabolism | 4-aminobutyraldehyde dehydrogenase (EC 1.2.1.19) |
| Amino Acids and Derivatives | Arginine; urea cycle, polyamines | Polyamine Metabolism | Spermidine Putrescine ABC transporter permease component PotB (TC 3.A.1.11.1) |
| Amino Acids and Derivatives | Arginine; urea cycle, polyamines | Polyamine Metabolism | Spermidine Putrescine ABC transporter permease component potC (TC_3.A.1.11.1) |
| Clustering-based subsystems | Clustering-based subsystems | CBSS-292415.3.peg.234 1 | Major facilitator superfamily (MFS) transporter |
| Clustering-based subsystems | no subcategory | Conserved gene cluster associated with Met-tRNA formyltransferase | 16S rRNA (cytosine(967)-C(5))-methyltransferase (EC 2.1.1.176) |
| Cofactors, Vitamins, Prosthetic Groups, Pigments | Biotin | Biotin biosynthesis | Long-chain-fatty-acid--CoA ligase (EC 6.2.1.3) |
| Cofactors, Vitamins, Prosthetic Groups, Pigments | Folate and pterines | 5-FCL-like protein | Butyryl-CoA dehydrogenase (EC 1.3.8.1) |
| Cofactors, Vitamins, Prosthetic Groups, Pigments | Tetrapyrroles | Cobalamin synthesis | Cobalamin synthase (EC 2.7.8.26) |
| DNA Metabolism | DNA repair | DNA repair, bacterial UvrD and related helicases | ATP-dependent DNA helicase UvrD/PcrA, proteobacterial paralog |
| DNA Metabolism | no subcategory | DNA structural proteins, bacterial | DNA-binding protein HBSu |
| Iron acquisition and metabolism | no subcategory | Hemin transport system | Outer membrane receptor proteins, mostly Fe transport |
| Membrane Transport | Cation transporters | Magnesium transport | Mg(2+) transport ATPase protein C |
| Membrane Transport | Cation transporters | Transport of Nickel and Cobalt | HoxN/HupN/NixA family nickel/cobalt transporter |
| Metabolism of Aromatic Compounds | Peripheral pathways for catabolism of aromatic compounds | Benzoate degradation | Benzoate 1,2-dioxygenase alpha subunit (EC 1.14.12.10) |
| Metabolism of Aromatic Compounds | Peripheral pathways for catabolism of aromatic compounds | Benzoate degradation | Benzoate 1,2-dioxygenase beta subunit (EC 1.14.12.10) |
| Metabolism of Aromatic Compounds | no subcategory | Aromatic Amin Catabolism | Nitrilotriacetate monooxygenase component B (EC 1.14.13.-) |
| Metabolism of Aromatic Compounds | no subcategory | Aromatic Amin Catabolism | Phenylacetaldehyde dehydrogenase (EC 1.2.1.39) |
| Phosphorus Metabolism | no subcategory | Phosphate metabolism | Soluble pyridine nucleotide transhydrogenase (EC 1.6.1.1) |
| Protein Metabolism | Protein biosynthesis | tRNA aminoacylation, Pro | tRNA proofreading protein STM4549 |
| Protein Metabolism | Protein processing and modification | G3E family of P-loop GTPases (metallocenter biosynthesis) | Urease accessory protein UreD |
| Protein Metabolism | Protein processing and modification | G3E family of P-loop GTPases (metallocenter biosynthesis) | Urease accessory protein UreE |
| Protein Metabolism | Protein processing and modification | G3E family of P-loop GTPases (metallocenter biosynthesis) | Urease accessory protein UreF |
| Protein Metabolism | Protein processing and modification | G3E family of P-loop GTPases (metallocenter biosynthesis) | Urease accessory protein UreG |
| Protein Metabolism | Protein processing and modification | G3E family of P-loop GTPases (metallocenter biosynthesis) | Urease alpha subunit (EC 3.5.1.5) |
| Protein Metabolism | Protein processing and modification | G3E family of P-loop GTPases (metallocenter biosynthesis) | Urease beta subunit (EC 3.5.1.5) |
| Protein Metabolism | Protein processing and modification | G3E family of P-loop GTPases (metallocenter biosynthesis) | Urease gamma subunit (EC 3.5.1.5) |
| Respiration | Electron accepting reactions | Anaerobic respiratory reductases | Vanillate O-demethylase oxidoreductase (EC 1.14.13.-) |
| Respiration | no subcategory | Carbon monoxide dehydrogenase maturation factors | Aerobic carbon monoxide dehydrogenase molybdenum cofactor insertion protein CoxF |
| RNA Metabolism | no subcategory | Group II intron-associated genes | Retron-type RNA-directed DNA polymerase (EC 2.7.7.49) |
| Stress Response | Oxidative stress | Oxidative stress | Redox-sensitive transcriptional activator SoxR |
| Sulfur Metabolism | Organic sulfur assimilation | Alkanesulfonate assimilation | Alkanesulfonate monooxygenase (EC 1.14.14.5) |
| Sulfur Metabolism | Organic sulfur assimilation | Alkanesulfonate assimilation | Alkanesulfonates ABC transporter ATP-binding protein |
| Sulfur Metabolism | Organic sulfur assimilation | Alkanesulfonate assimilation | Alkanesulfonates-binding protein |

**Table 10:** Identification of RAST Subsystems exclusive to the ATCC 33445 strain of *Acetobacter pasteurianus*.

| Category | Subcategory | Subsystem | Role |
| --- | --- | --- | --- |
| Amino Acids and Derivatives | Arginine; urea cycle, polyamines | Arginine Biosynthesis -- gjo | Acetylornithine deacetylase (EC 3.5.1.16) |
| Amino Acids and Derivatives | Lysine, threonine, methionine, and cysteine | Threonine degradation | Threonine dehydrogenase and related Zn-dependent dehydrogenases |
| Carbohydrates | no subcategory | Conserved cluster around inner membrane protein gene yghQ, probably involved in polysaccharide biosynthesis | Conserved hypothetical TPR repeat protein, clustered with yghQ |
| Carbohydrates | no subcategory | Conserved cluster around inner membrane protein gene yghQ, probably involved in polysaccharide biosynthesis | Conserved protein YghT, with nucleoside triphosphate hydrolase domain |
| Cell Wall and Capsule | Capsular and extracellular polysacchrides | Rhamnose containing glycans | capsular polysaccharide biosynthesis protein |
| Clustering-based subsystems | no subcategory | CBSS-316273.3.peg.2378 | FIG006126: DNA helicase, restriction/modification system component YeeB |
| Clustering-based subsystems | no subcategory | CBSS-316273.3.peg.2378 | FIG045374: Type II restriction enzyme, methylase subunit YeeA |
| Clustering- | no | CBSS-316273.3.peg.2378 | YeeC-like protein |

| based<br>subsystems | subcategory |  |  |
| --- | --- | --- | --- |
| Cofactors,<br>Vitamins,<br>Prosthetic<br>Groups,<br>Pigments | Riboflavin,<br>FMN, FAD | Riboflavin, FMN and FAD<br>metabolism | 3,4-dihydroxy-2-butanone 4-phosphate synthase (EC 4.1.99.12) |
| Cofactors,<br>Vitamins,<br>Prosthetic<br>Groups,<br>Pigments | Tetrapyrroles | Cobalamin synthesis | Cobalamin biosynthesis protein CbiG |
| DNA<br>Metabolism | no<br>subcategory | DNA structural proteins,<br>bacterial | DNA-binding protein HU |
| DNA<br>Metabolism | no<br>subcategory | Restriction-Modification<br>System | Type I restriction-modification system,<br>specificity subunit S (EC 3.1.21.3) |
| Fatty Acids,<br>Lipids, and<br>Isoprenoids | Fatty acids | Fatty Acid Biosynthesis FASII | Enoyl-[acyl-carrier-protein] reductase<br>[NADPH] (EC 1.3.1.10) |
| Membrane<br>Transport | no<br>subcategory | Ton and Tol transport<br>systems | TolA protein |
| Miscellaneous | no<br>subcategory | Broadly distributed proteins<br>not in subsystems | Putative oxidoreductase YncB |
| Nitrogen<br>Metabolism | Denitrification | Denitrifying reductase gene<br>clusters | Respiratory nitrate reductase alpha<br>chain (EC 1.7.99.4) |
| Nitrogen<br>Metabolism | Denitrification | Denitrifying reductase gene<br>clusters | Respiratory nitrate reductase beta<br>chain (EC 1.7.99.4) |
| Nitrogen<br>Metabolism | Denitrification | Denitrifying reductase gene<br>clusters | Respiratory nitrate reductase delta<br>chain (EC 1.7.99.4) |
| Nitrogen<br>Metabolism | Denitrification | Denitrifying reductase gene<br>clusters | Respiratory nitrate reductase gamma<br>chain (EC 1.7.99.4) |
| Nitrogen<br>Metabolism | no<br>subcategory | Nitrate and nitrite<br>ammonification | Assimilatory nitrate reductase large<br>subunit (EC:1.7.99.4) |
| Nitrogen<br>Metabolism | no<br>subcategory | Nitrate and nitrite<br>ammonification | Nitrate ABC transporter, nitrate-<br>binding protein |
| Nitrogen<br>Metabolism | no<br>subcategory | Nitrate and nitrite<br>ammonification | Nitrate/nitrite transporter |
| Nitrogen<br>Metabolism | no<br>subcategory | Nitrate and nitrite<br>ammonification | Nitrite reductase [NAD(P)H] large<br>subunit (EC 1.7.1.4) |
| Nitrogen<br>Metabolism | no<br>subcategory | Nitrate and nitrite<br>ammonification | Response regulator NaST |
| Phages,<br>Prophages,<br>Transposable<br>elements,<br>Plasmids | Phages,<br>Prophages | Phage tail proteins | Phage tail length tape-measure<br>protein |
| Phages,<br>Prophages, | Phages,<br>Prophages | Phage tail proteins | Phage tail tube protein |
| Transposable elements, Plasmids |  |  |  |
| Phages, Prophages, Transposable elements, Plasmids | Phages, Prophages | Phage tail proteins | Phage tail/DNA circulation protein |
| Protein Metabolism | Protein biosynthesis | tRNAs | tRNA-Ser-CGA |
| Protein Metabolism | Protein biosynthesis | tRNAs | tRNA-Ser-GGA |
| Protein Metabolism | Protein processing and modification | N-linked Glycosylation in Bacteria | N-linked glycosylation glycosyltransferase PglG |
| Regulation and Cell signaling | Programmed Cell Death and Toxin-antitoxin Systems | Phd-Doc, YdcE-YdcD toxin-antitoxin (programmed cell death) systems | Prevent host death protein, Phd antitoxin |
| Regulation and Cell signaling | Programmed Cell Death and Toxin-antitoxin Systems | Phd-Doc, YdcE-YdcD toxin-antitoxin (programmed cell death) systems | Programmed cell death antitoxin MazE like |
| Regulation and Cell signaling | Programmed Cell Death and Toxin-antitoxin Systems | Phd-Doc, YdcE-YdcD toxin-antitoxin (programmed cell death) systems | Programmed cell death toxin MazF like |
| Regulation and Cell signaling | Programmed Cell Death and Toxin-antitoxin Systems | Toxin-antitoxin replicon stabilization systems | RelE/StbE replicon stabilization toxin |
| RNA Metabolism | RNA processing and modification | ATP-dependent RNA helicases, bacterial | Cold-shock DEAD-box protein A |
| Stress Response | no subcategory | Flavohaemoglobin | Nitric-oxide reductase (EC 1.7.99.7), quinol-dependent |
| Sulfur Metabolism | Organic sulfur assimilation | Alkanesulfonate assimilation | probable dibenzothiophene desulfurization enzyme |
| Virulence, Disease and Defense | Resistance to antibiotics and toxic compounds | Arsenic resistance | Arsenic efflux pump protein |

| Virulence,<br>Disease and<br>Defense | Resistance to<br>antibiotics<br>and toxic<br>compounds | Arsenic resistance | Arsenic resistance protein ArsH |
| --- | --- | --- | --- |

In this comparative analysis, we noted four subsystems exclusive to the AD and NBCR 101655 genomes that may explain their ability to survive passage through the fly intestine. Both strains encode polyamine metabolism factors that are frequently associated with cellular growth and survival, and have established roles in the formation of biofilms (43). The association-competent genomes also encode factors necessary for the conversion of urea to ammonium and carbon dioxide. A similar system operates in *Helicobacter pylori* where it raises the gastric pH to generate a more hospitable environment for microbial survival (44). We also detected the redox-sensitive transcriptional activator SoxR in both association-competent genomes. SoxR promotes microbial survival within the intestine by countering the antibacterial actions of host-derived superoxide anions (45). Finally, we detected several genes that contribute to organic sulfur assimilation in association-competent genomes. These gene products allow *A. pasteurianus* AD and NBRC 101655 to use alkanesulfonates as a source of sulfur during sulfate or cysteine starvation and may provide both strains a competitive advantage in the intestine if sulfur is limiting.

The association-incompetent ATCC 33445 strain also encodes products that contribute to generation of ammonia. However, the ATCC 33445 strain relies on respiratory nitrate reductase and nitrite reductase to generate ammonia, as well as assimilatory nitrate reductase to access nitrate for metabolic growth. This represents an entirely different strategy to use nitrogen as a fuel for metabolic energy and growth. Thus, it is feasible that the ATCC strain fails to grow within the adult intestine due to unsatisfied nitrogen requirements. We also observed two toxin-antitoxin systems unique to the association-incompetent ATCC 33445 genome – an addiction module toxin that ensures propagation of plasmids to progeny cells (46), and a MazE/MazF type toxin-antitoxin (47). The MazE/MazF system induces programmed cell death in prokaryotic cells in response to stressful environments. It is intriguing to speculate that the adult intestine activates this response, thereby inducing microbial death and preventing ATCC 33445 survival within the gut.

### Lactobacillus plantarum

#### General Genomic Features

We used the environmental ATCC 14917 strain as a reference for our comparative analyses of *L. plantarum* genomes. *L. plantarum* ATCC 14917 was isolated from pickled cabbage and we confirmed viable association of this strain with the adult *Drosophila* intestine. We compared the ATCC 14917 strain to four *Drosophila*-associated genomes – WJL, DMCS_001, DF and KP. WJL and DMCS_001 were isolated from *Drosophila* raised in geographically separate labs (48, 49). We isolated the KP strain from the intestines of our lab-raised wild-type strain, and the DF strain from an isofemale wild *Drosophila melanogaster* line that we captured in Edmonton, Canada in the summer of 2014. The DF and KP genomes encode one chromosome and three closely related plasmids each (Figure 4). While all five genomes are closely related, genome-to-genome distance calculators suggest a greater degree of identity among the *Drosophila*-associated KP, DF, WJL and DMCS_001 strains (Table 11). In general, the environmental genome is smaller than the *Drosophila*-associated genomes (Figure 5A), encodes fewer RNAs and coding sequences, and contains fewer phage-associated proteins (Table 11).

**Figure 4:**
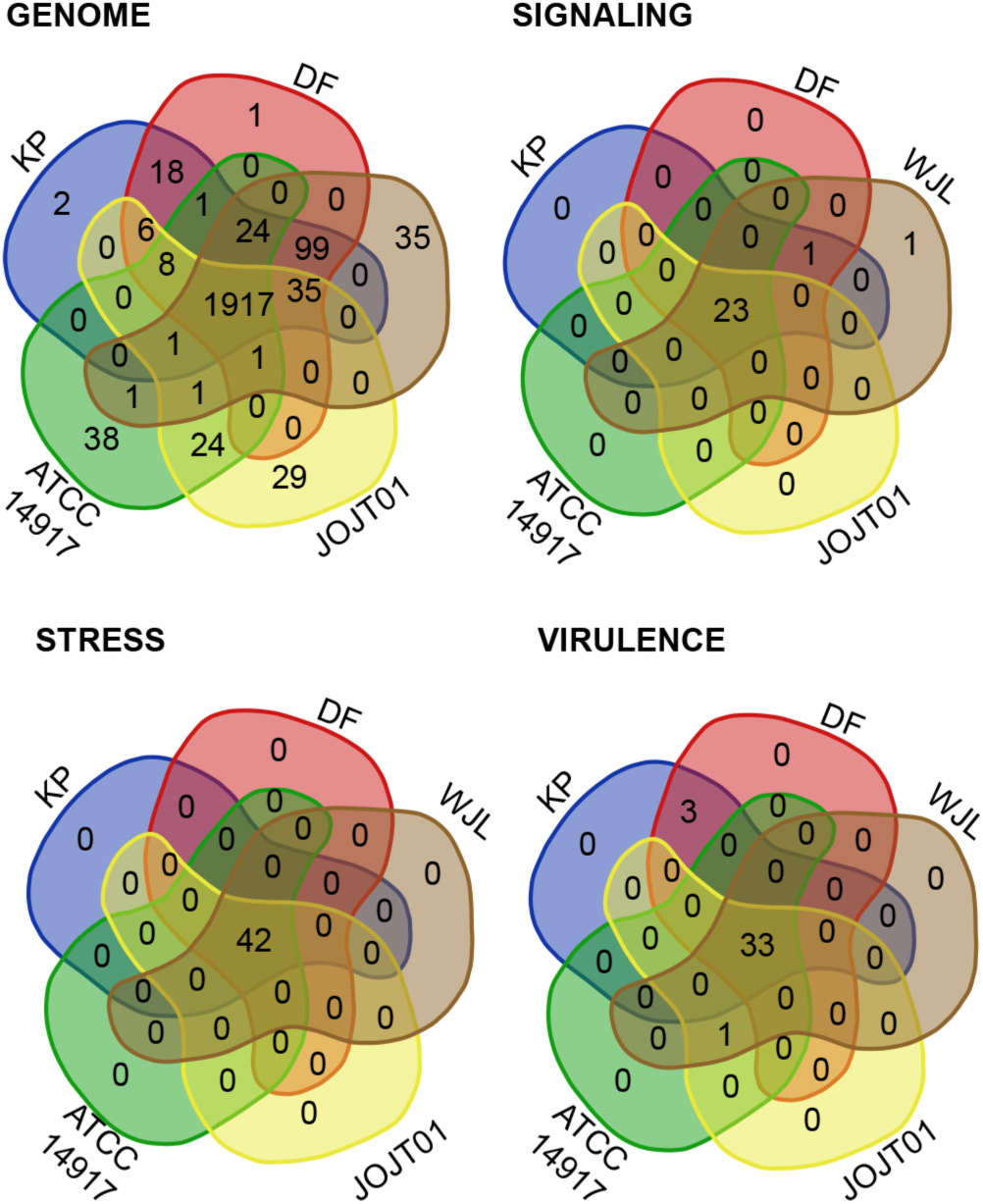
Distribution of unique gene functions in the genomes of *L. plantarum* strains KP, DF, WJL, JOJTOI and ATCC 14917. All data are based on gene function annotations within RAST and exclude gene products with unknown functions.

**Figure 5:**
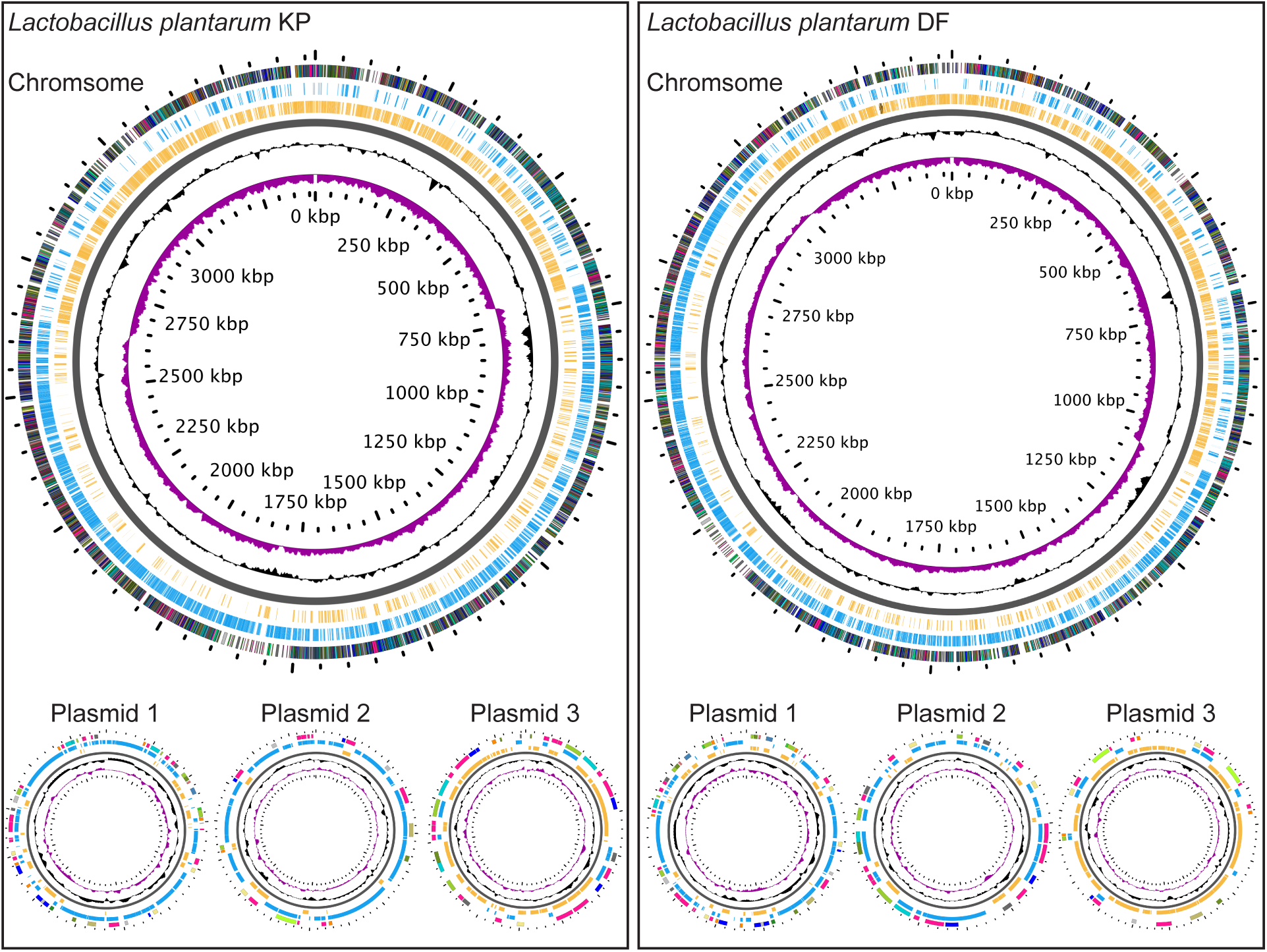
Illustrations of the genomes for *L. plantarum* strains KP and DF. GC skew is indicated in purple, and GC content is indicated in black. All positive strand ORFs are shown in blue, and negative strand ORFs are shown in yellow.

**Table 11:** Details on *Lactobacillus plantarum* genomes described in this study

|  | *L. plant* KP | *L. plant* DF | *L. plant* WJL | *L. plant* JOJT01.1 | *L. plant* ATCC 14917 |
| --- | --- | --- | --- | --- | --- |
| Number of contigs or scaffolds | 4 | 4 | 102 | 83 | 36 |
| Genome to genome distance (Prob. DDH >= 70%) | 100 | 98.22 | 97.74 | 97.04 | 96.99 |
| Genome size | 3,692,742 | 3,697,306 | 3,477,495 | 3,194,687 | 3,198,761 |
| CG content (%) | 44 | 44.5 | 44.2 | 44.5 | 44.5 |
| Number of RNAs | 104 | 104 | 85 | 65 | 65 |
| Predicted CDS | 3569 | 3574 | 3365 | 3063 | 3061 |
| Assigned function | 2403 | 2400 | 2344 | 2227 | 2246 |
| Uncharacterized | 9 | 8 | 9 | 9 | 9 |
| Conserved hypothetical | 13 | 12 | 10 | 10 | 11 |
| Unknown function | 72 | 72 | 74 | 69 | 70 |
| Hypothetical | 884 | 894 | 797 | 694 | 668 |
| Phage-associated proteins | 180 | 188 | 131 | 54 | 57 |

#### Environmental Response Factors

Contrary to our expectations, we did not observe pronounced differences between the genomes of environmental or *Drosophila*-associated strains in terms of modifiers of signal transduction, stress responses or virulence (Figure 5 B-D, and supplementary tables 13–15). Likewise, there is not a notable enrichment of two-component systems, transcription factors or other DNA binding proteins in *Drosophila-associated* strains of *L. plantarum* (Table 12, supplementary tables 16–18). In general, these findings suggest that *Drosophila*-associated strains of *L. plantarum* are not easily distinguished on the basis of functions that sense and respond to specific environmental cues.

**Table 12:** Organization of regulatory proteins within *Lactobacillus plantarum* genomes described in this study

|  | *L. plant* KP | *L. plant* DF | *L. plant* WJL | *L. plant* JOJT01.1 | *L. plant* ATCC 14917 |
| --- | --- | --- | --- | --- | --- |
| Two-component systems | 27 | 27 | 29 | 25 | 25 |
| Transcription Factors | 252 | 252 | 244 | 230 | 230 |
| Other DNA binding proteins | 20 | 20 | 18 | 13 | 13 |

**Table 13:** Prophage and CRISPR identification in *Lactobacillus plantarum* genomes described in this study

|  | *L. plant* KP | *L. plant* DF | *L. plant* WJL | *L. plant* JOJT01.1 | *L. plant* ATCC 14917 |
| --- | --- | --- | --- | --- | --- |
| Number of Prophages | 5 | 6 | 4 | 2 | 2 |
| Identity of Prophages | 2 X NC_00435<br>(*Lactobacillus* phage) | 2 X NC_00435<br>(*Lactobacillus* phage) | NC_00435<br>(*Lactobacillus* phage) | NC_009812<br>(*Lactobacillus* phage) | NC_005355<br>(*Lactobacillus* phage) |
|  | NC_023559<br>(*Oenococcus* phage) | NC_023559<br>(*Oenococcus* phage) | NC_023560<br>(*Oenococcus* phage) | NC_019489<br>(*Lactobacillus* phage) | NC_019489<br>(*Lactobacillus* phage) |
|  | 2 X NC_019489<br>(*Lactobacillus* phage) | 3 X NC_019489<br>(*Lactobacillus* phage) | NC_019489<br>(*Lactobacillus* phage) |  |  |
|  |  |  | NC_024384<br>(*Lactobacillus* phage) |  |  |
| Number of CRISPRs | 0 | 0 | 0 | 0 | 0 |
| CRISPR targets | 0 | 0 | 0 | 0 | 0 |

**Table 14:**
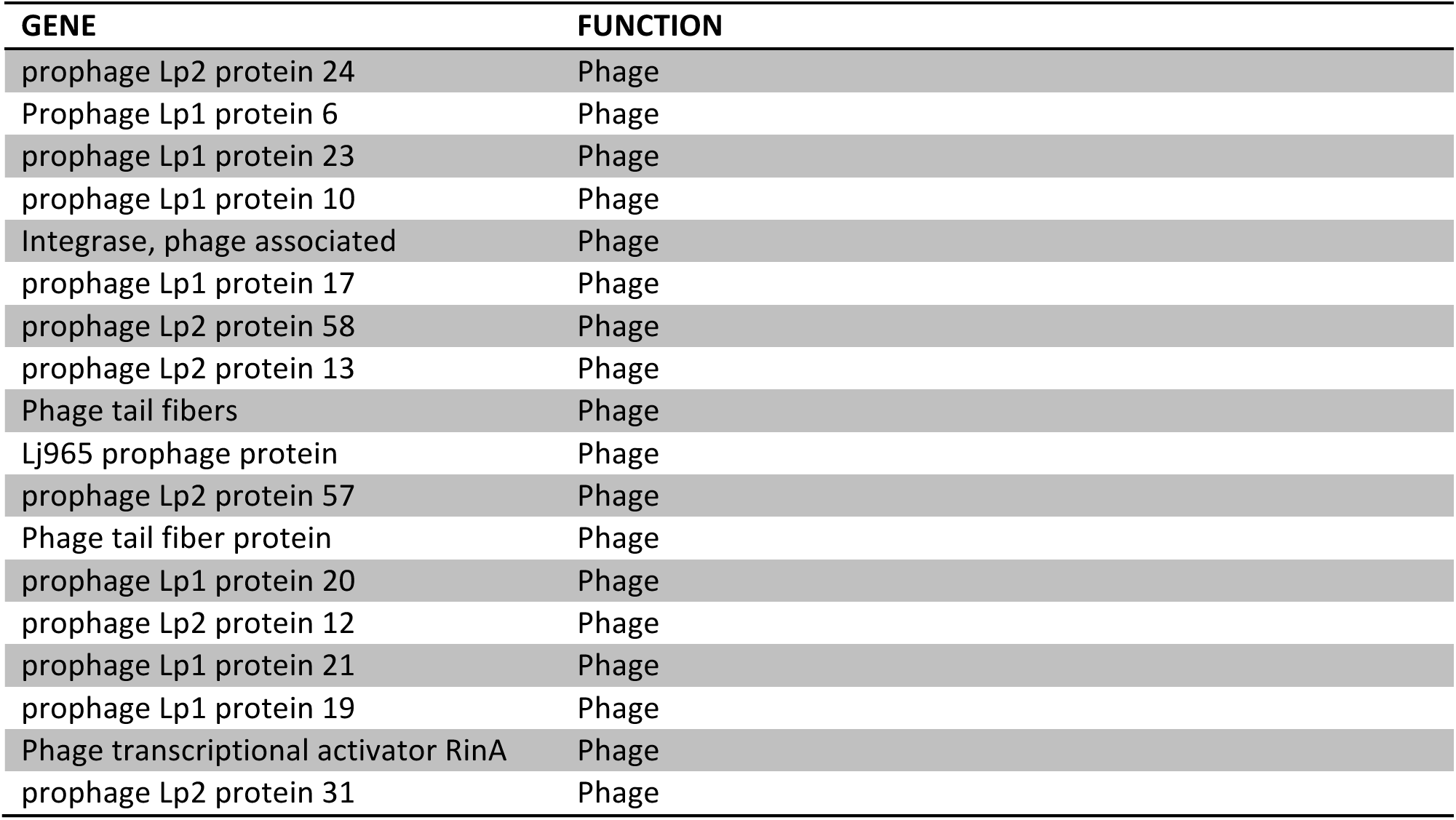
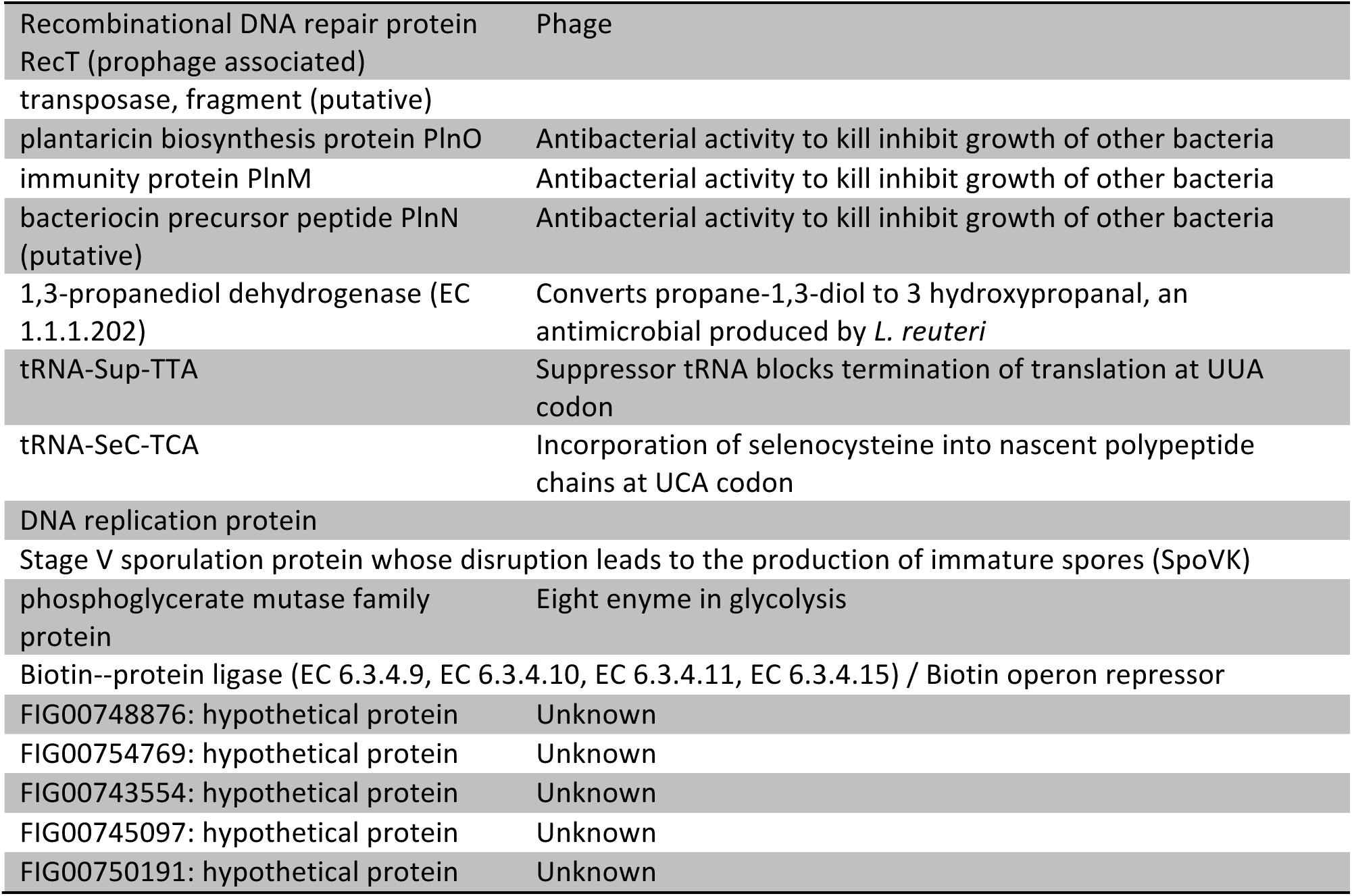
Identification of RAST Subsystems unique to *Drosophila*-associated strains of *Lactobacillus plantarum*.

#### Prophages and CRISPR Responses

Similar to our observations with *L. brevis* genomes, we observed a greater number of intact prophage genomes in *Drosophila-associated L. plantarum* strains than in the environmental strain (Table 13). Of all *Lactobacillus* genomes studied, we detected an average of four intact prophage genomes in *Lactobacillus* strains isolated from flies, and a maximum of two prophage genomes in environmental strains (Figure 6). The main difference between the *Drosophila*-associated *brevis* and *plantarum* strains is that the *plantarum* strains do not appear to encode CRISPR-dependent anti-phage defenses within their genomes.

**Figure 6:**
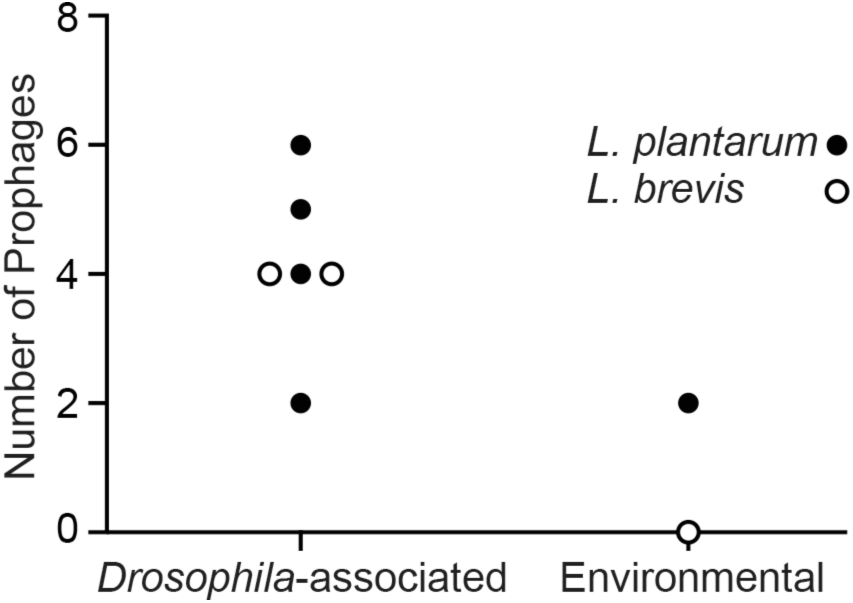
Numbers of intact prophage genomes associated with *L. brevis* genomes (open circles) and *L. plantarum* genomes (closed circles) described in this study.

#### Function-Based Comparisons of Commensal and Reference Strains of *Lactobacillus plantarum*

We then used the whole-genome sequence data for environmental and *Drosophila*-associated strains to prepare an *L. plantarum* pangenome (Figure 7). Visual inspection indicated that three of the *Drosophila-associated* genomes (DF, KP and WJL) display similar organization, while the DMCS_001 genome appears more akin to the environmental genome. When we looked for factors exclusively common to the DF, KP and WJL genomes we detected several gene products that derive from prophages. However, we also detected functional elements that are consistent with support for microbial growth in the intestinal lumen. This includes access to methionine, an essential amino acid in *L. plantarum;* enzymes for the regulation of sialic acid, a comparatively rare bacterial molecule that is typically found in prokaryotes that live in association with deuterostomes (50); and a regulator of the biosynthesis of lipoteichoic acid, a regulator of cell wall autolysis in gram-positive bacteria (51, 52). Strikingly, we also detected uridine phosphorylase exclusively in the genomes of the *Drosophila*-associated DF, KP and WJL strains. Uridine phosphorylase catalyzes conversion of uridine to uracil, a secreted metabolite that causes microbial-dependent inflammation in the intestine of adult *Drosophila* (53). Combined, these genes constitute a comparatively small number of molecular functions that may favor the propagation and establishment of *L. plantarum* populations in the fly intestine.

**Figure 7:**
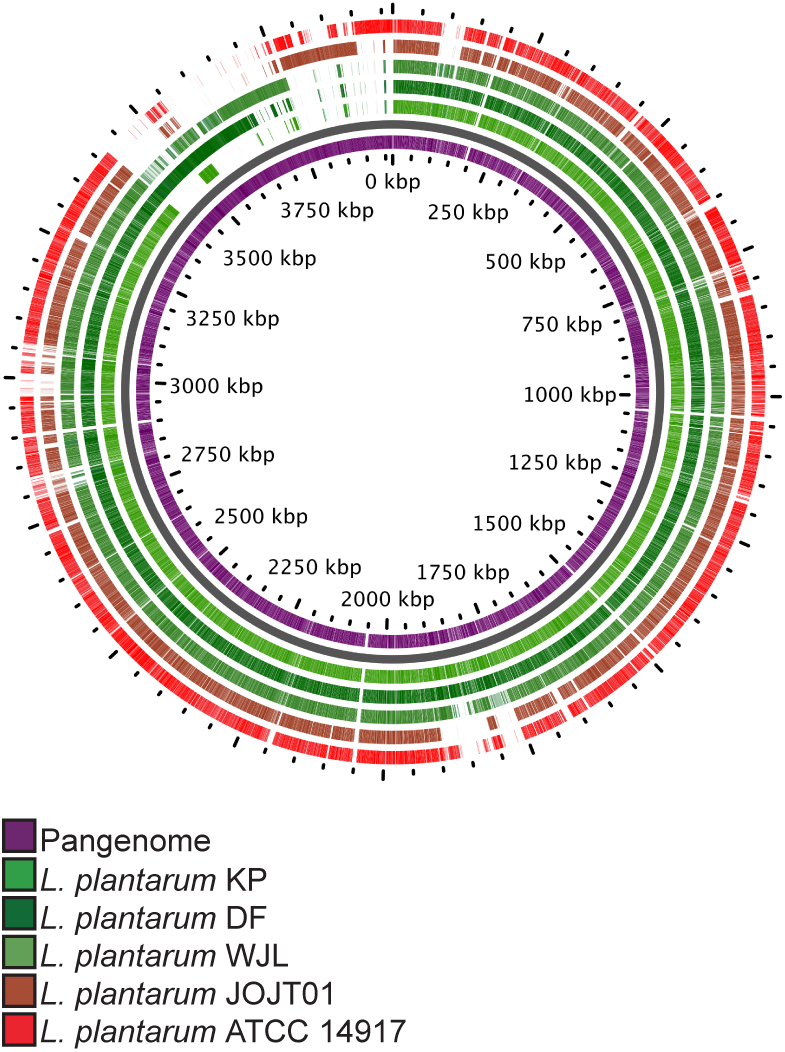
The inner wheel shows a pangenome for all *L. plantarum* strains described in this study, with the corresponding genomes aligned in outer wheels.

Finally, we examined the thirty-five genes exclusively observed in the genomes of DF, KP, WJL and DMCS_001. The majority (nineteen) were prophage genes, and an additional five were hypothetical proteins of unknown function. Rather strikingly, several of the remaining genes encode products that actively suppress the growth of competing microbes. These include the *PlnMNO* operon that encodes a bacteriocin and cognate immunity protein (54), and 1,3-propanediol dehydrogenase, an enzyme that converts propane-1,3,-diol to 3-hydroxypropanal. 3-hydroxypropanal is also known as reuterin, a *Lactobacillus reuteri* metabolite that exerts broad-spectrum microbicidal effects on intestinal microbes (55). Combined the actions of *PlnMNO* and reuterin give *Drosophila*-associated *L. plantarum* a competitive advantage against other microbes within the gut lumen.

## DISCUSSION

The last decade witnessed a proliferation of elegant studies that uncovered critical host responses to microbial factors in the *Drosophila* intestine (Reviewed in (13)). Bacterial cues promote larval growth (32, 33), direct innate immune responses (56, 57), orchestrate the proliferation of intestinal stem cells (58, 59), and regulate the uptake and storage of nutrients (36). Despite the importance of the intestinal microflora for *Drosophila* health and development, we know remarkably little about the biochemical events that permit bacterial survival within the hostile terrain of a fly intestine. A recent study identified microbial metabolism and stress response pathways that mediate interactions between intestinal bacterial and their *Drosophila* host (49). In this study, we examined the genomes of *Drosophila*-associated strains of *L. brevis, L. plantarum*, and *A pasteurianus*. We were particularly interested in the identification of candidate signal transduction, virulence and defense factors that permit bacterial survival within the intestines of adult flies. To this end, we compared fly-associated genomes to environmental strains of the same species. For each species, we observed a small number of genetic pathways that were exclusive to the *Drosophila*-associated genomes. Many of the *Drosophila*-associated pathways encode products with established roles in host-microbe interactions, suggesting these products facilitate association of *Drosophila* with the individual strains.

### Lactobacilli

For our studies of *Lactobacillus* genomes, we prepared whole-genome sequences of *L. brevis* or *L. plantarum* strains that we isolated from lab-raised wild-type flies, and an *L. plantarum* strain that we isolated from a wild *Drosophila*. These genomes formed the cornerstones of a comparative study that included three previously reported *Drosophila*-associated genomes (37, 48, 49), and the genomes of environmental strains that successfully associate with the intestines of our wild-type *Drosophila*. In this manner, we identified bacterial functions that are unique to *Drosophila*-associated genomes of *L. brevis* and *L. plantarum*. The functions fall into four broad categories – antibacterial, structural, metabolic, and phage-related.

The most striking feature common to all four *Drosophila*-associated *L. plantarum* genomes was the presence of broad-spectrum bactericidal factors. For example, the DF, KP, WJL and DMCS_001 genomes all encode a complete *PnIMNO* operon, which encodes a bacteriocin and a corresponding immunity protein (54). Bacteriocins are produced by many lactic acid bacteria to kill neighboring bacteria, while the immunity protein protects *L. plantarum* from collateral damage (60). In addition, the *Drosophila*-associated *L. plantarum* genomes encode the enzymatic capacity to generate 3-hydroxypropanal/reuterin, a bacterial toxin expressed by *L. reuteri* in the gut to suppress the growth of other commensals. Combined, these bactericidal molecules counter the growth of competing bacteria inside a *Drosophila* host, and favor expansion of *L. plantarum*. The competitive advantages conferred by the *PnlMNO* operon and 3-hydroxypropanal may explain why *L. plantarum* is frequently reported in studies that characterize the intestinal microflora of *Drosophila*.

The *Drosophila*-associated genomes of *L. plantarum* and *L. brevis* also encode structural components that stabilize associations with their fly host. In particular, we detected metabolic pathways for modifications to cell wells that permit host-microbe interactions and biofilm formation. These include the construction of exopolysaccharides by *L. brevis* and the regulation of sialic acid by *L. plantarum*. Sialic acid is a comparatively rare microbial metabolite, but has been observed on microbes that associate with deuterostomes. Bacteria use sialic acid as a nutrient, but they also use it to mask detection by host immune responses. While the role of sialic acid in *L. plantarum* association with *Drosophila* requires further investigation, we feel that these elements merit consideration as host-microbe interaction factors.

The *Drosophila*-associated genomes of *L. plantarum* and *brevis* also includes gene products that address nutritional requirements. These functions include access to limited resources such as methionine by *L. plantarum*, and utilization of citrate as an energy source by *L. brevis*. We were particularly struck by the presence of pectin metabolism factors within the genomes of *Drosophila*-associated strains of *L. brevis*. Pectin is an excellent source of carbon for bacteria that grow on plants. However, bacterial utilization of pectin accelerates the ripening and decay of the same plants (61). Thus, *Drosophila-associated L. brevis* genomes express factors that contribute to the decay of organic substrates. This is particularly notable, as *Drosophila* preferably consumes decayed matter as a source of nutrients. The ability of *L. brevis* to generate meals for their *Drosophila* host provides a possible explanation for the fact that *Drosophila* frequently associate with *L. brevis*, even though intestinal bacteria do not appear to establish stable populations within the guts of adult *Drosophila*. As *L. brevis* generates palatable meals for fly hosts, we speculate that their chances of association with flies in the wild are greater than those for many other bacteria. This host-microbe relationship is similar to a proposed mechanism for association of *Erwinia carotovora* with *Drosophila* in the wild (62).

The final difference we noted between environmental and *Drosophila*-associated *Lactobacillus* genomes was an accumulation of temperate prophage genomes throughout *Drosophila*-associated *Lactobacilli*. Intestinal stresses such as high levels of reactive oxygen species are known to trigger lysogenic induction of temperate prophages (63). Thus, it is feasible that bacterial strains that pass though the fly intestine will release and integrate greater numbers of lytic prophages, explaining the increased numbers of prophage genomes in *Lactobacillus* strains that associate with adult *Drosophila*.

### Acetobacter

In this study, we report the first genome of a *Drosophila*-associated strain of *A. pasteurianus*, and identified an *A. pasteurianus* strain that fails to survive passage through the fly gut. A comparison of association-competent and association-incompetent genomes is limited by the fact that only one *Drosophila*-associated genome is available for study. Nonetheless, our identification of an environmental *A. pasteurianus* strain that fails to associate with *Drosophila* yields a comparatively short list of candidate functions that regulate association of *Acteobacter* with adult *Drosophila*. This list includes gene products that process nitrogen, and gene products that directly control the induction of cell death in *A. pasteurianus*. To our knowledge, ours is the first study to identify a set of bacterial factors that regulate association of an *Acetobacter* with *Drosophila*. We believe that future characterization of mutations in the respective gene products will identify the bacterial factors that control viability of *A. pasteurianus* in the adult intestine. This approach has considerable value given the relationship between *Acetobacter* and *Drosophila* development.

In summary, our comparative study of bacterial genomes uncovers a number of genetic signatures of association with *Drosophila*. As many of the gene products have established roles in host-microbe interactions, we propose that these genes include factors that promote the frequent association of *Drosophila* with *Lactobacillus* and *Acetobacter* strains. Future characterization of mutations in the individual products will reveal the relationship between the individual factors and host physiology.

## CONCLUSION

The intestinal microflora influences a number of physiological outcomes in their animal host. Microbial factors shape the establishment and execution of immune responses, influence the availability of critical nutrients and often determine the progression of debilitating inflammatory illnesses. Studies in *Drosophila melanogaster* have provided valuable insights into the host cellular machinery that coordinates responses to microbial presence. However, we know remarkably little about the microbial factors that permit long-term association with the host. In this study, we characterized the genomes of common *Drosophila*-associated microbes and identified putative bacterial products that favor association with *Drosophila*. We believe that products constitute valuable starting points to examine the microbial half of the host-microbe relationship.

## MATERIALS AND METHODS

### *Drosophila* husbandry

All *Drosophila* assays were performed with virgin *w^1118^* male and female flies raised on standard corn-meal medium (Nutri-Fly Bloomington Formulation, Genesee Scientific) in a humidified incubator at 29°C. To generate germ-free flies, we transferred freshly eclosed (0 – 16 hr old) adult flies to standard medium that we supplemented with an antibiotic (100 μg/ml ampicillin, 50 μg/ml vancomycin, 100 μg/ml neomycin and 100 μg/ml metronidazole dissolved in ethanol). This cocktail has been described previously (22). To generate gnotobiotic flies, we raised adult flies on the antibiotic cocktail for five days, starved flies for two hours, and transferred the flies to a vial containing an autoclaved fly vial cotton plug soaked with the respective bacteria resuspended in a sterile-filtered solution of 10% sucrose in PBS. We fed the flies the bacterial meal for 16 hours and then transferred the flies to vials of freshly autoclaved food. To confirm association, we periodically plated fly homogenates on bacterial medium selective for *Acetobacter* or *Lactobacilli*.

### Bacterial Isolation and Sequencing

We plated homogenates of 15 day old adult *Drosophila* on Acetobacter and MRS culture plates. We found that *L. brevis* colonies are easily distinguished from *L. plantarum* colonies on MRS-agar medium. We isolated individual colonies of *A. pasteurianus, L. brevis* and the KP strain of *L. plantarum* and grew them statically at 29°C in liquid MRS (L. *brevis* and *L. plantarum)*, or shaking in liquid (A. *pasteurianus)*. The DF strain of *L. plantarum* was isolated from a wild, mated isofemale *Drosophila melanogaster* captured on a rotting strawberry in the kitchen of EF in Edmonton, Canada. Bacterial DNA was isolated with the Microbial DNA Isolation kit from MO BIO Laboratories Inc. (catalog number: 12224–250) according to their instructions. The genomes of *L. plantarum* strains DF and KP were sequenced and assembled at the McGill University and Génome Québec Innovation Centre on the PacBio platform. The genomes of *A. pasteurianus* (strain AD) and *L. brevis* (strain EF) were sequenced at The Applied Genomics Core of the University of Alberta. For the latter genomes, we prepared Nextera XT libraries from the isolated micribial DNA according to Illumina’s protocol and sequenced the libraries with using the V3-600 cycle Kit (Illumina). Whole genome sequences were then assembled using Lasergene software (DNASTAR). This Whole Genome Shotgun project for *L. brevis* EF has been deposited at DDBJ/EMBL/GenBank under the accession LPXV00000000. The version described in this paper is version LPXV01000000. The Whole Genome Shotgun project for *A. pasteurianus* AD has been deposited at DDBJ/EMBL/GenBank under the accession LPWU00000000. The version described in this paper is version LPWU01000000. Chromosome1 of *L. plantarum* KP has been deposited at DDBJ/EMBL/GenBank under the accession CP013749 and plasmids 1–3 for the same strain have been deposited under the accession numbers CP013750, CP013751 and CP013752, respectively. Chromosome1 of *L. plantarum* DF has been deposited at DDBJ/EMBL/GenBank under the accession CP013753 and plasmids 1–3 for the same strain have been deposited under the accession numbers CP013754, CP013755 and CP013756, respectively.

### Genome Assembly and Annotation

For each sequencing project, we confirmed the individual species with the SpeciesFinder 1.2 algorithm (64) and calculated genome to genome distances with the genome to genome distance calculator of the Leibniz Institute DSMZ-German Collection of Microorganisms and Cell Cultures (65). We annotated each genome with RAST (66), identified intact prophage genomes with the PHAST server (67) and identified CRISPR arrays with CRISPRFinder (68). We used the CRISPRTarget algorithm to predict CRISP targets for the individual CRISPR arrays (69). To identify regulatory proteins within the respective genomes, we used the P2RP identifier (70). We used the GView tool to generate graphical representations of bacterial genomes (71).

## ACKNOWLEDGEMENTS

This research was funded by a grant from the Canadian Institutes of Health Research to EF (MOP 77746). Next Generation Sequencing services were performed by The Applied Genomics Core (TAGC) at the Faculty of Medicine & Dentistry, University of Alberta.

